# Truncated WT1 protein isoform expression is increased in MCF-7 cells in a long-term estrogen depletion

**DOI:** 10.1101/802439

**Authors:** Saavedra-Alonso Santiago, Zapata-Benavides Pablo, Mendoza-Gamboa Edgar, Chavez-Escamilla Ana Karina, Arellano-Rodríguez Mariela, Rodriguez-Padilla Cristina

## Abstract

**Background:** The WT1 gene codes for a transcription factor that presents several protein isoforms with diverse biological properties, capable of positively and negatively regulating genes involved in proliferation, differentiation, and apoptosis. WT1 protein is overexpressed in more than 90% of breast cancer, however, its role during tumor progression is still unknown.

**Methodology:** In this work were analyzed the expression of WT1 isoforms (36-38 kDa and 52-54 kDa, and 17 AA (+/−) and KTS (+/−)) in breast cancer cells. On the other hand, with the purpose of mimicking the process of switch from a hormone-dependent to a hormone-independent neoplasm, an assay was performed using the MCF-7 cells cultured in long-term estrogen depletion (MCF-7 LTED cells) to determine the WT1 protein isoforms expression by western blot and RT-PCR, and Her2/neu and Estrogen receptor (ER) expression by quantitative RT-PCR assay. Growth kinetics and sensitivity to tamoxifen were performed in the MCF-7 LTED cells by trypan blue exclusion.

**Results:** The western blot shows the presence of the 52-54 kDa WT1 isoform in the ER (+) breast cancer cells, but not in the ER (−) cells. The 36-38 kDa WT1 isoform was detected in all the breast cancer cell lines analyzed. Using specific primers was found that 17 AA (+) / KTS (−) WT1 isoform was the most frequent in four breast cancer cell lines. During the sampling of the MCF-7 cells in estrogen depletion, an increase in the short-term of 52-54 kDa WT1 isoform was observed and this was kept until week 13, thereafter, its expression was absent; alternately, the 36-38 kDa WT1 isoform was observed from week 1 and it remained constant until week 27. MCF-7 LTED cells growth kinetic decreased 1.4 folds and were not sensitive to tamoxifen antiproliferative effect (p ≤ 0.05). Finally, were observed an increase of expression of ER and Her2/neu in the MCF-7 LTED cells.

**Conclusions:** The 36-38 kDa WT1 isoform expression occurs during the modifications of the hormonal environment, suggesting that it may be playing an important role in its adaptation and tumor progression.

## Introduction

The Wilms’ tumor gene (WT1) consists of 10 exons that encode a zinc finger transcription factor involved in the genitourinary development during embryogenesis (1, 2). Recently studies describe WT1 protein as an oncogene because is enveloped in cancer regulating genes responsible for cell growth, apoptosis and tumoral angiogenesis such as cyclin D1, Bcl-2, Bcl-xL, BFL1, c-myc, VEGF (3–8). The WT1 protein is conformed by an N-terminal protein region contains domain involved in transcriptional regulation, self-association, and RNA recognition (9–11), and a C-terminal protein region that contains a DNA/RNA binding domain and consist in four zinc-finger that can bind to GC-rich sequences (12–14).

WT1 gene shows two alternative splicing sites, identified as exon 5 or 17AA and other that occurs in the exon 9 and identified as KTS (Lys-Thr-Ser) (15). The different isoforms are referred to as A) isoform that lacks both 17 amino acid and KTS inserts; B) isoform contains the 17 amino acid insert but lacks KTS insert; C) isoform lacks the 17 amino acid insert but contains KTS insert; and D) isoform contains both inserts (16, 17). Moreover, has three sites of initiation of translation that produces three isoforms of different molecular weight (62-64 kDa, 52-54 kDa and 36-38 kDa) with different biological properties (18, 19). The 62-64 kDa WT1 protein isoform is not essential for normal development and reproduction in mice (*20*) and, the 36-38 kDa WT1 protein isoform has more oncogenic potential than WT1 52-54 kDa protein isoform in leukemia cells (20, 21).

It is reported that WT1 appears to have a growth regulatory role in many solid cancers including lung (22), colon (23), pancreatic (24), breast (25), and gastric (26), and no solid cancer as leukemia (27), and lymphoma (28). In breast cancer, Silberstein et al., 1997 initially considered to WT1 as a tumor suppressor gene, because its overexpression was found in healthy tissue and not in breast cancer tissue (29), however; subsequent work by Loeb et al., 2001, reported that WT1 expression has been found in 90% of samples breast cancer (25). Miyoshi et al., 2002, correlated high levels of WT1 mRNA with poor prognosis of survival rate in breast cancer patients (30), and Oji et al., 2004, demonstrated that the wild-type WT1 gene plays an important role in the tumorigenesis of breast cancer (31).

Currently, it is not fully known what is the interaction that WT1 presents with the tumor markers used in the characterization of breast cancer, such as the Estrogen Receptor, Progesterone Receptor, and Her2/neu. Studies in breast cancer lines have demonstrated WT1 expression involved with modulation of tumoral markers expression. Zapata-Benavides et al., 2002, found that WT1 protein decreased around 60% in MCF-7 cells cultured without estrogen stimulus, and WT1 expression was restored after 17β-estradiol incubation (32). Another work-related to tumor markers in breast cancer was performed by Tuna et al., 2005, where they observe that the signaling pathway of Her2/neu through Akt it is possible to regulate the levels of WT1 expression, concluding that the WT1 protein plays a vital role in the regulation of cell cycle progression and apoptosis (33). Accordingly, due to the previous studies, we hypothesize that overexpression of WT1 and the change of isoforms allow WT1 to be present at different stages of breast cancer progression.

In this paper, were analyzed the expression of WT1 isoforms, ER and Her2/neu in a long-term estrogen depletion *in vitro* model, to mimic the malignant progression of breast cancer switch from estrogen-dependent to estrogen-independent growth.

## Methodology

### Cellular culture

To initial assays, MCF-7, BT-474, T47D, SKBR-3, MDA-MB-231, MDA-MB-453, and BT-20 breast cancer cells were obtained from the ATCC (American Type Culture Collection, Manassas VA). All cell lines were grown in DMEM/F12 medium supplemented with 10% fetal bovine serum (FBS) and incubated in 95% air with 5% CO_2_ at 37°C.

### Quantitative RT-PCR

The total RNA of tissue was isolated using 1 mL of reagent Trizol (Life Technologies, Gaithersburg, MD) according to the manufacturer’s instructions. The DNAc was performed using 5 μg of total RNA, Superscript II and oligo (dT_12-18_) under the condition at 42°C for 90 min, followed by heating at 70°C for 10 minutes. Each reaction of PCR real-time was made using 2 μL of DNAc. To Her2/neu was used forward primer 5’-GAGGCACCCAGCTCTTTGA-3’, reverse primer 5’-CGGGTCTCCATTGTCTAGCA-3’ and probe Fam-5’-CCAGGGCATAGTTGTCC-3’NFQ, and to Estrogen Receptor was used forward primer CGACATGCTGCTGGCTACA, reverse primer 5’-ACTCCTCTCCCTGCAGATTCAT-3’ and probe Fam-5’-CATGCGGAACCGAGATGA-3’NFQ. Human β-actin primers set manufactured by Applied Biosystems were used as endogen control. For each reaction was used Universal PCR Master Mix manufactured by Roche Branchburg, New Jersey USA. The protocol was performed for 40 cycles at 94°C for 30 seconds and 64°C for 30 seconds. The assay of relative expression was conducted using the Livak method (34).

### PCR to detect spliced isoforms of 17AA / KTS

Ratios of isoform spliced of WT1 gene, 17AA (−) / KTS (−), 17AA (−) /KTS (+), 17AA (+) / KTS (−) and 17AA (+) / KTS (+) to total WT1 transcripts were obtained by PCR according of Yosuke Oji et al., 2002 (22). The sequences of primers were as follows: F2 5’-GACCTGGAATCAGATGAACTTAG-3 and R2 5-GAGAACTTTCGCTGACAAGTT-3’ to determine the ratio of 17AA (+) / (−); F3 5’-GTGTGAAACCATTCCAGTGTA-3’ and R3 5’-TTCTGACAACTTGGCCACCG-3’ to determine the ratio of KTS (+) / (−). PCR amplification was performed with 35 cycles at denaturalization 94°C for 60 sec, alignment at 56°C for 60 sec, and extension of 72°C for 90 sec. All PCR products were analyzed on 10% acrylamide gels and visualized with ethidium bromide under UV light.

### Western blot to detect WT1 isoforms protein

Total protein collection was using reagent Trizol according to the manufacturer’s instructions. Protein samples (50 μg) were electrophoresed on 12% SDS–polyacrylamide gels and transferred to nitrocellulose membranes. Immunodetection to detect the 52-54 kDa WT1 protein isoform was performed using WT-1 (6F-H12, NH-terminal binding) monoclonal antibody obtained from DAKO (Carpinteria, CA) and to detect all isoforms was used WT1 (C19, COOH-terminal binding) polyclonal antibody obtained from Santa Cruz Biotechnology, β-actin monoclonal antibody obtained from Sigma Chemical (St Louis, MO) and anti-mouse and anti-rabbit antibodies conjugated with horseradish peroxidase were purchased from Bio-ad. Protein bands were visualized by enhanced chemiluminescence using Roche Lumi-Light Western blotting substrate.

### Long-Term Estrogen Depletion assay

MCF-7 cells were grown in DMEM/F12 culture medium supplemented with 10% FBS until 60% confluence, thereafter the cells were washed twice with sterile buffer PBS to eliminate the phenol red, and culture medium was replaced with DMEM/F12 phenol red-free supplemented with 10% of charcoal-dextran treated FBS manufactured by HyClone (Road Logan Utah, USA), the time of sampling were made according to cell growth.

### Growth kinetics

The growth was performed seeding 2.5 ×10^5^ MCF-7 cells in a normal condition and MCF-7 LTED cells (27 weeks) per well in six wells plates using respective culture medium than were harvested at 24 and 72 hours and quantified by trypan blue exclusion staining. Each count was done by triplicate and the initial number of cells were considered as 1.

### Assays of MCF-7 LTED cells treated with tamoxifen

MCF-7 (3×10^3^) cells in a normal condition and MCF-7 LTED cells (27 weeks) were seeded per well in a plate of 96 wells, after 24 hours of incubation, was added the 1.25, 2.5, 5 and 10μM of tamoxifen concentrations, each treatment was carried out by triplicate including control of untreated cells, the plates were then incubated for 24 hours, then were analyzed using the MTT assay. The reagent MTT (3-(4, 5-dimethylthiazol-2-yl)-2, 5- diphenyl tetrazolium bromide was purchased from Sigma-Aldrich. MTT solution was prepared to 5 μg/mL in PBS buffer, then 20 μL of MTT solution was added per well, and then the plate was incubated at 37°C. Finally, the medium was removed, and 200 μL of DMSO was added to solubilize the formazan salt. The plate was analyzed in a microplate reader (Microplate Autoreader EL311, BioTek Instruments Inc., Winooski, Virginia, USA) at 570 nm to a determinate optical density (OD).

### Statistical analysis

Significance of different treatments was determined by analysis of variance and Student’s t-test. Differences were considered significant at p ≤ 0.05 using SPSS software, version 13 (SPSS, Inc., Chicago, IL, USA). All data were expressed as the mean ± the standard errors. P<0.05 was considered to indicate a statistically significant difference. To calculate the IC_50_ was used the AAT Bioquest, Inc. (2019, September 02). Quest Graph™ IC_50_ Calculator (https://www.aatbio.com/tools/ic50-calculator).

## Results

### WT1 protein isoforms expression in breast cancer cell lines

Initially, the characterization of the expression of the WT1 isoforms was carried out in breast cancer cell lines MCF-7, BT-474, T47D, SKBR-3, MDA-MB-231, MDA-MB-453, and BT-20. WT1 protein expression was detected in all these cell lines, and with different isoform patterns. The 52-54 kDa WT1 isoform was expressed only in the positive estrogen receptor cell lines (MCF-7, BT-474, and T47D), but was not expressed in the negative estrogen receptor cell lines (tumor markers status is showed in **Table 1)**: SKBR-3, MDA-MB-231, MDA-MB-453, and BT-20 (**Figure 1A**). Moreover, the 36-38 kDa WT1 isoform was expressed in all these breast cancer cells. To determine alternative splicing 17AA/KTS, were analyzed four of the seven cell lines. The MCF-7 cells expressed four isoforms and the BT-474, SKBR-3, MDA-MB-231 cells only expressed the 17AA + / KTS − isoform (**Figure 1B**).

**Table 1.** Tumor markers status of breast cancer cell lines

| Cell lines | Tumor markers |  |  |
| --- | --- | --- | --- |
|  | Estrogen receptor | Progesterone receptor | Her2/neu |
| MCF-7 | + | - | - |
| BT-474 | + | + | + |
| T47D | + | + | - |
| SKBR-3 | - | - | + |
| MDA-MB-231 | - | - | - |
| MDA-MB-453 | - | - | Low |
| BT-20 | - | - | - |

Present (+); ausent (-); Her2/neu, human epidermal growth receptor 2

**Figure 1.**
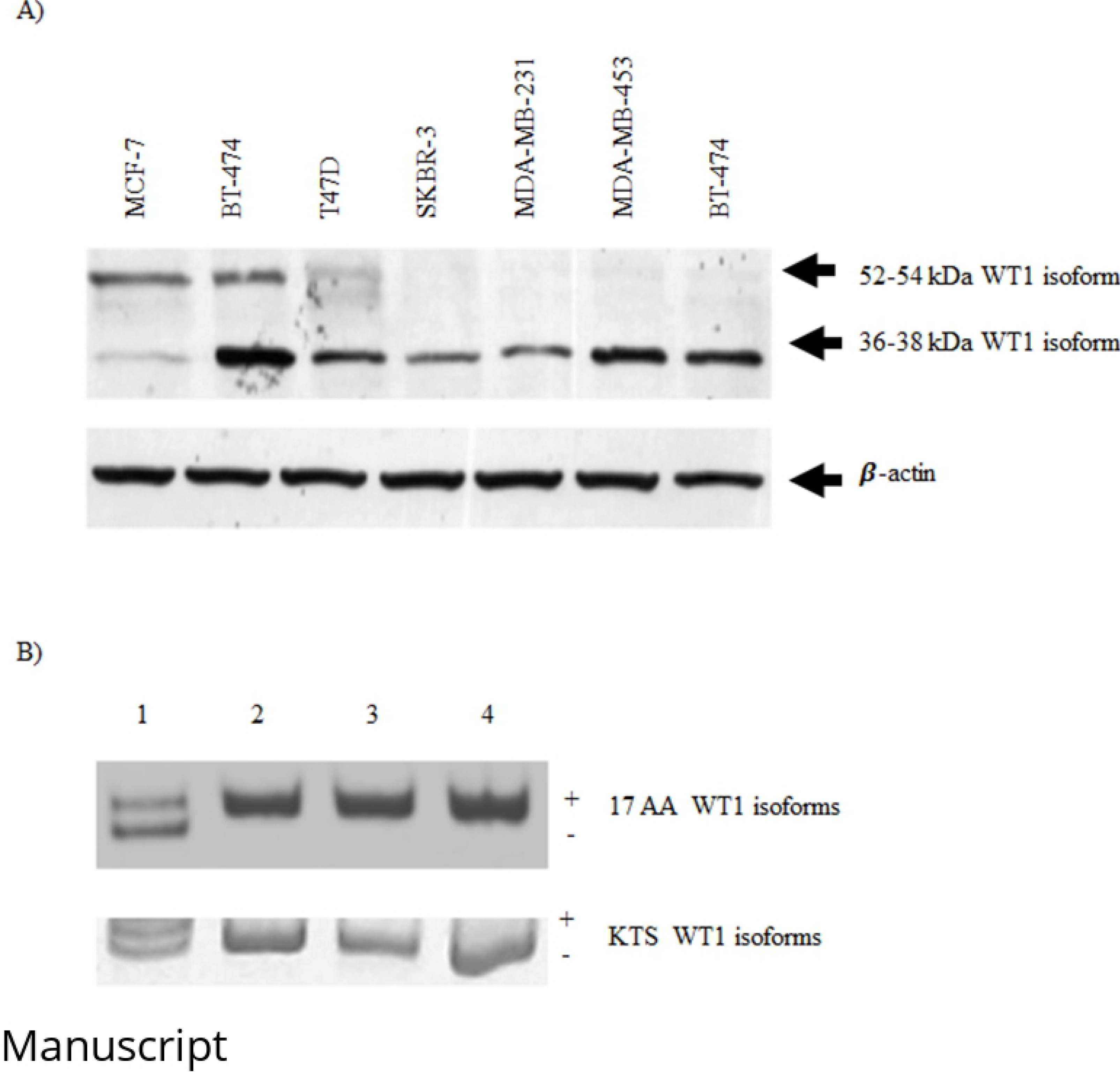
Expression of the WT1 isoforms in breast cancer cell lines. A) The 52-54 kDa and 36-38 kDa WT1 isoforms were analyzed in breast cancer cell lines. The expression of β-actin was included as an endogenous control. B) Analysis of alternative splicing 17AA / KTS in breast cancer cell lines (lane 1 MCF-7; lane 2 BT474; lane 3 SKBR-3; and lane 4 MDA-MB-231).

### MCF-7 cells in short-term estrogen depletion increases the expression of 52-54 kDa WT1 protein isoform

To analyze the effects of estrogen depletion in the WT1 expression, several assays were carried out in the MCF-7 positive estrogen receptor cells. The MCF-7 cells cultured under normal conditions was changed to medium to conditions of estrogen depletion for 24 and 48 hours, then a western blot was performed. The expression of the 36-38 kDa WT1 protein isoform was not affected, however, the expression of the 52-54 kDa WT1 protein isoform was significantly increased (**Figure 2A**).

**Figure 2.**
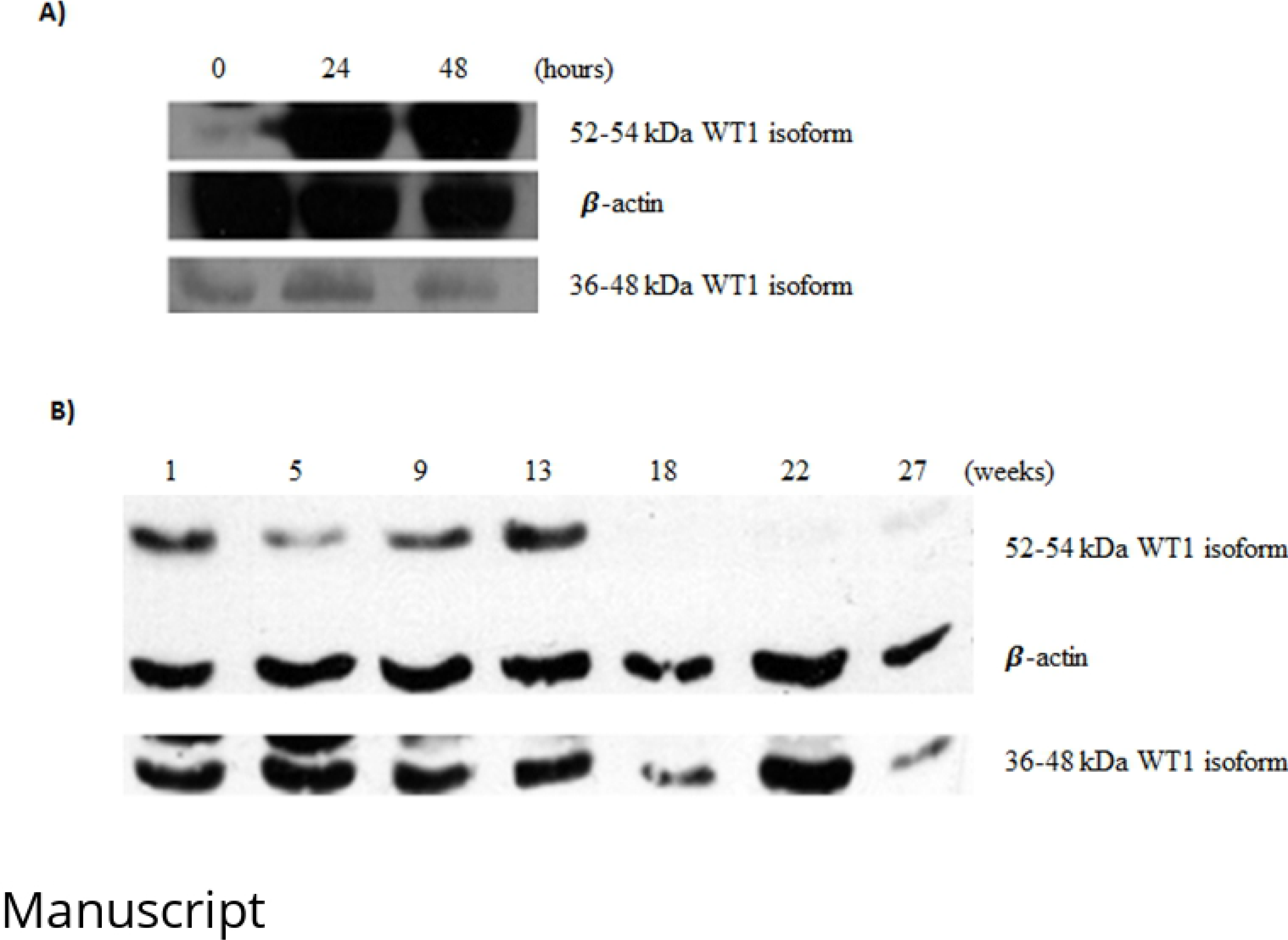
Expression of WT1 in the MCF-7 cells cultured in estrogen depletion. A) The 36-38 kDa and 52-54 kDa WT1 isoforms expression in MCF-7 cells in depletion of estrogen in a short-term (hours). B) The 36-38 kDa and 52-54 kDa WT1 isoforms expression in MCF-7 cells in depletion of estrogen in a long-term estrogen depletion (weeks). In both figures, β-actin expression was used as an endogenous control.

### MCF-7 cells in long-term estrogen depletion increases the expression of 52-54 kDa and 36-38 kDa WT1 protein isoforms

Subsequently, samples of MCF-7 cells were collected during several weeks in long-term estrogen depletion (MCF-7 LTED cells) and were analyzed to determine the expression of the WT1 isoforms by western blot. The expression of 52-54 kDa WT1 protein isoforms significantly increased until the 13th week, however, disappears through the 18th week. The expression of 36-38 kDa WT1 protein isoforms significantly increased during all the 27 weeks (**Figure 2B**).

### The low proliferation of MCF-7 cells cultured in long-term estrogen depletion

Growth kinetic was performed to compare the MCF-7 cells cultured in the medium under normal conditions against the MCF-7 LTED cells cultured for 27 weeks in culture medium with estrogen depletion. The assay was performed using the culture medium corresponding to each of the estrogen conditions. MCF-7 LTED cellular proliferation was significantly decreased (1.4 folds) (value p ≤ 0.05) in comparison with the MCF-7 cells cultivated under normal conditions (**Figure 3A**).

**Figure 3.**
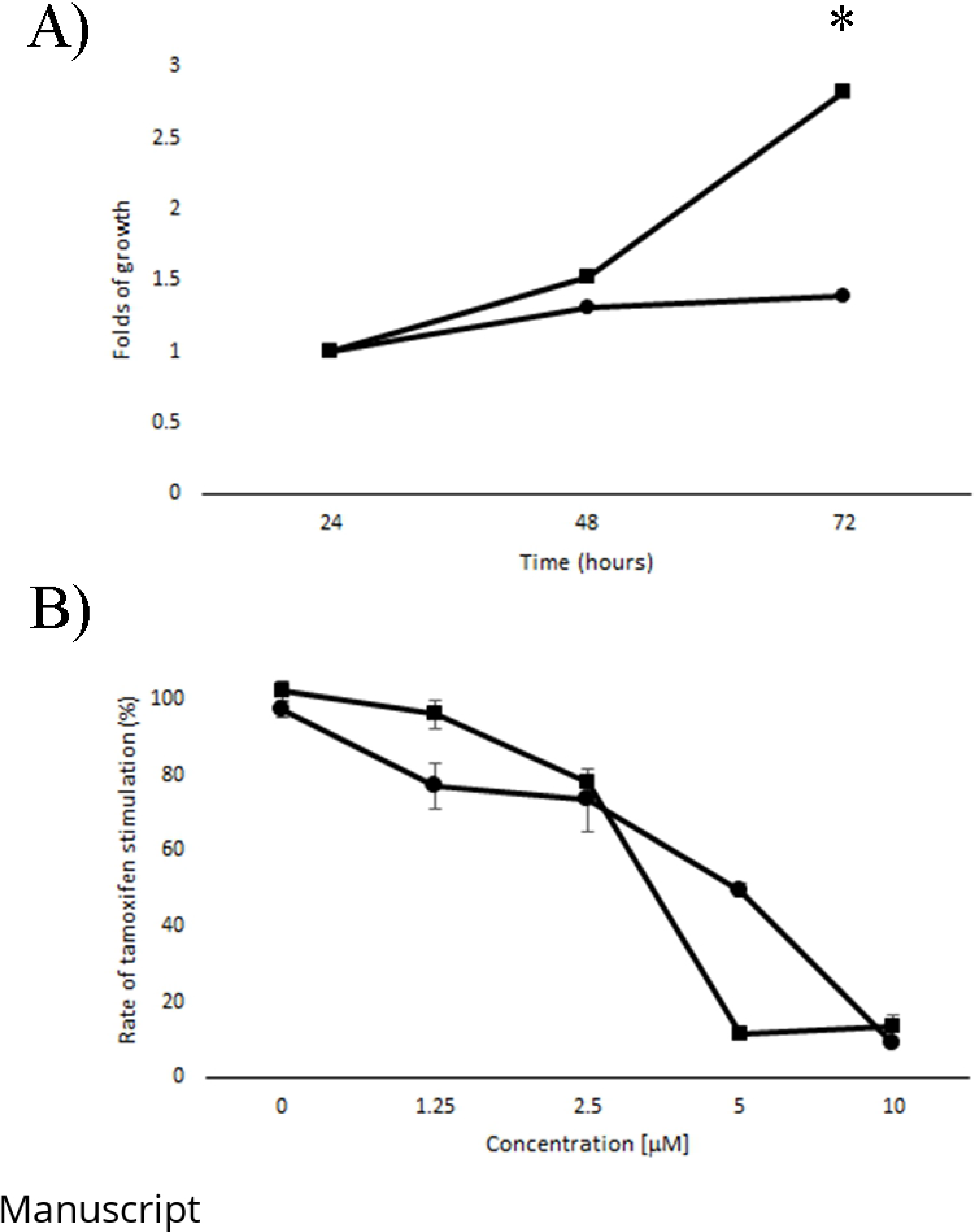
Growth kinetics and Tamoxifen effect on MCF-7 LTED cells. A) The growth of MCF-7 cells with 27 weeks in estrogen depletion was compared to MCF-7 cells cultured in a normal condition. Count of cells was analyzed to 24 and 48 hours. Each assay was carried out by triplicate including the standard error. B) The sensibility of tamoxifen using the following concentration 1.25μM, 2.5μM, 5μM, and 10μM was analyzed on MCF-7 LTED cells (⚫) and MCF-7 cells in a normal condition (⬛). All data are the mean ± SD of three independent experiments (*p< 0.05).

### Effect of tamoxifen on MCF-7 LTED cells

Subsequently, a tamoxifen sensitivity test was performed on the MCF-7 LTED cells (27 weeks) compared to the MCF-7 cells cultured under normal conditions. MCF-7 LTED cells were less sensitive IC_50_ = 4.935 mM of inhibition effect to tamoxifen in comparison to the MCF-7 cells in normal conditions IC_50_ = 3.278 mM (p ≤ 0.05) (**Figure 3B**). The MCF-7 LTED cells were much less sensitive to the anti-proliferative effect of tamoxifen.

### Analysis of Her2/neu and Estrogen Receptor Expression in MCF-7 LTED cells

To observe the behavior of the molecular markers Her2/neu and Estrogen Receptor a quantitative RT-PCR analysis was performed using samples of MCF-7 LTED cells taken in several times. The mRNA expression of Her2/neu and ER, significantly increased in MCF-7 LTED cells, that started from week 22 until week 53, was reported an increase of 2.98 (SD ± 0,43) and 2.59 (SD ± 0,99) expression levels of the markers Her2/neu and Estrogen Receptor, respectively (**Figure 4**).

**Figure 4.**
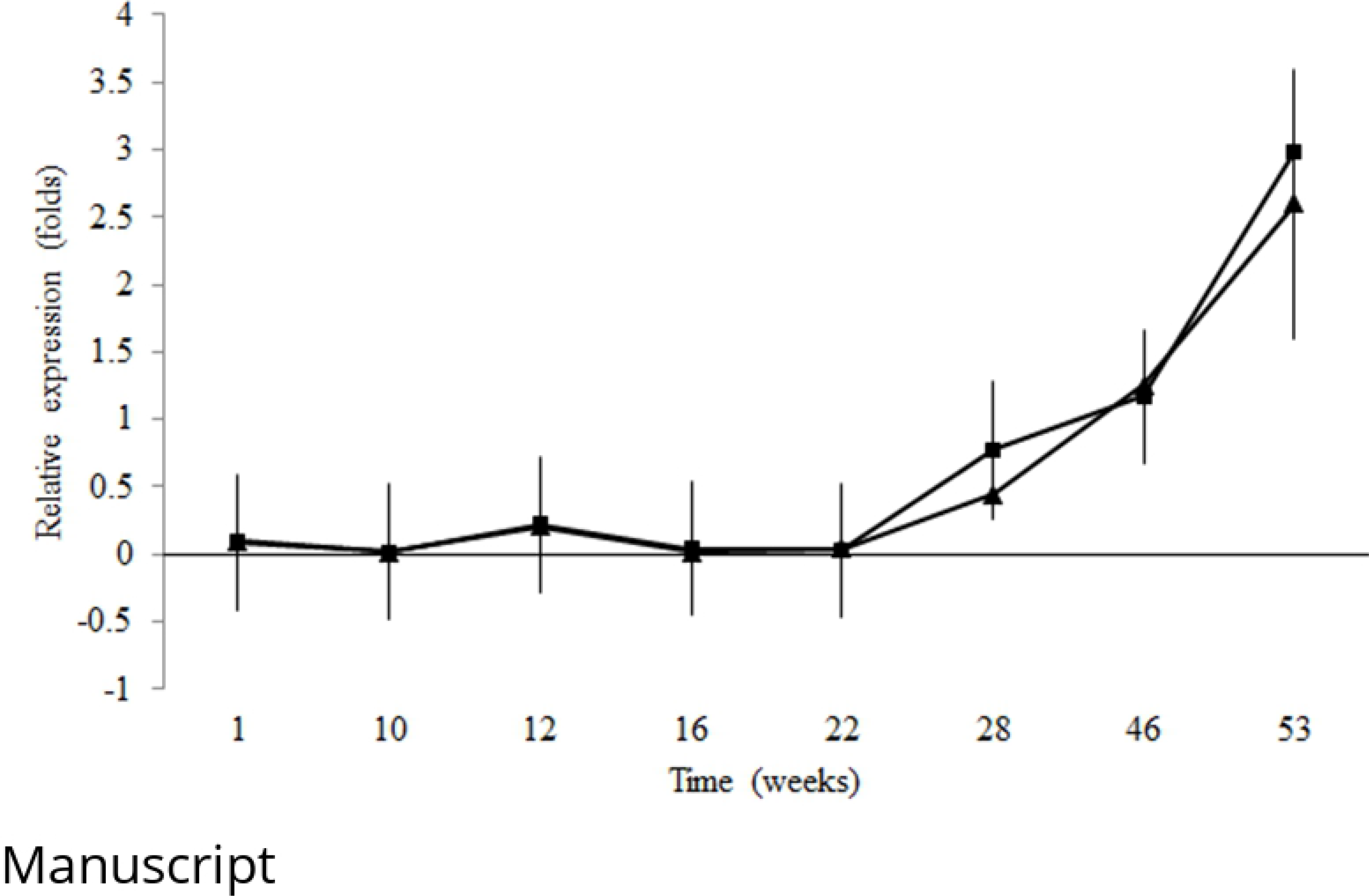
Expression of tumor markers Her2/neu and Estrogen Receptor in MCF-7 LTED cells by quantitative RT-PCR assay. The graph shows the relative expression of Her2 / neu (⬛) and ER (▲). All the experiments were done by triplicate and including β-actin as an endogenous control.

## Discussion

Breast cancer worldwide represents a major health problem, due to its high incidence and high mortality that occupies the second cause of cancer death in general, and the first cause of cancer death in women (35). The molecular classification of breast cancer is performed to the tumor markers Progesterone receptor, Estrogen receptor and Her2/neu. Estrogen receptor and Her2/neu are the most important tumor markers in breast cancer because their presence dictates the type of therapy to be followed and the prognosis of the disease (36, 37). The critical step in malignant progression of breast cancer is the switch from estrogen-dependent to estrogen-independent growth (38). During the change of hormonal dependence, the activity of Her2/neu is increased, relating it to a high metastatic potential (39).

High expression of WT1 wild-type in breast cancer has been reported in > 90% (25, 30), and high levels of the WT1 expression have been inversely associated with patient survival. The presence of WT1 is essential for breast cancer cell growth (32), however, it is still unclear what role it plays during the progression of breast cancer. WT1 can act as a tumor suppressor gene or as an oncogene, and this may be due to the presence of various isoforms which have different biological properties (40).

The four isoforms are expressed in primary human solid cancers, including lung cancer (30), HNSCC (41) sarcoma (42), breast cancer (43), and human primary leukemia (44). However, the functions of each of the four WT1 isoforms in cancer cells remain unclear (45). In the present study, were analyzed the WT1 isoforms present in the breast cancer cell lines. In our results, were observed the presence of the 52-54 kDa WT1 isoform in Estrogen Receptor-positive cell lines and this isoform was not detected in Estrogen Receptor-negative cell lines. The 36-38 kDa WT1 isoform was present in all these cell lines analyzed. The biological function of 36-38 kDa WT1 isoform is not clear yet. The 36-38 kDa WT1 isoform lacks the first 128 amino acids in the amino-terminal region, which generates a loss in the dimerization domain and a loss of the repression domain. Its presence has been determined in cell lines and specimens of the Wilms tumor where it has been reported that its transactivating function is 1.5 levels higher than the 52-54 kDa WT1 isoform KTS (−) (19).

In order to the alternative splicing of WT1, we observe that MCF-7 cells presented the four isoforms (17 AA (+/−) / KTS (+/−)) and the other cell lines only expressed the 17AA (+) / KTS (−) WT1 isoform. The 17 AA (+) WT1 isoforms are involved in cell proliferation, apoptosis, cancer development and protects cells against etoposide-induced apoptosis (46, 47). Ectopic overexpression of 17 AA (+)/KTS (+) and 17 AA (+)/KTS (−) WT1 isoforms in MCF-7 cells reduces pro-apoptotic BAK and caspase-7 proteins, and p53 mRNA levels (48). Burwell et al., 2007, in their work was associates the specific presence of 17 AA (+) / KTS (−) WT1 isoform with reduction in cell proliferation and with promoted the appearance of highly organized acinar cellular aggregates and 17AA (+) / KTS (+) WT1 isoform with caused an epithelial-mesenchymal transition and a redistribution of E-cadherin (43). Other work in vitro, analyze the 17AA (−) / KTS (−) WT1 isoform which is layers of modulation of expression of cytoskeletal regulatory proteins such as α-actinin 1, cofilin and gelsolin, allowing cancer cells to acquire a most aggressive phenotype (49).

Other works have associated the 17AA (+) / KTS (+) WT1 isoform with the differentiation block but cell proliferation is induced in 32D cl3 myeloid progenitor cells (50), and normal myeloid cells in response to granulocyte-CSF (51). Tuna M et al., 2017, showed that the treatment of MCF-7 cells with insulin-like growth factor I (IGF-I) increases WT1 protein expression by 77% in especially the 17 AA (+) / KTS (−) WT1 isoform (16). One of the possible hypotheses of the multiple activities of the WT1 isoforms may be due to protein-protein interactions in region 17 AA. Several publications have described the interactions of WT1 with proteins such as PAR-4 and p53, therefore, having to consider in future studies the presence of proteins that interact with the isoforms of WT1 (52–54).

Subsequently, were made a model cultivate the MCF-7 cells under conditions of estrogen depletion, to mimic the process that occurs naturally with menopause and the use of aromatase inhibitors as hormonal therapy in breast cancer. In our results analyzed in estrogen depletion trial, were shown an increase in the expression of the 52-54 kDa WT1 isoform at 24 hours and this expression was observed during the 13th week, however its expression disappears later. On the other hand, the expression of the 36-38 kDa WT1 isoform is not affected in the short-term trial, however, its expression was observed from the first week and is mostly constant until week 27th. We hypothesize that the increase in the expression of 36-38 kDa WT1 isoform is involved in the change during estrogen-depletion which may be associated with the loss of the repression domain of WT1 which would offset the decrease of the stimulus in the mediated proliferation by estrogens. The cell proliferation of cells cultured in estrogen depletion showed a significant decrease, in addition to presenting a slight insensitivity to Tamoxifen’s antiproliferative stimulation.

Then, we analyzed the quantitative expression of ER and Her2/neu and in both cases a gradual increase is observed at the time of cultivation of the MCF-7 cells in estrogen depletion increases, coinciding with the results of Lei Wang et al., 2010, they observe an estrogenic independence in MCF-7 cells through the mitogen-activated protein kinases (MAPK) pathway (38, 40).

Lei Wang et al., 2010, observed in a high passage model MCF-7 cells (MCF-7H) exhibited estrogen-independent and anti-estrogen insensitive growth MCF7H cells expressed high levels of Her2/neu, EGFR and ER-α in breast cancer cells (38), which would explain the increase in ER and Her2/neu transcripts in the cells treated in estrogen depletion. In general, our results show the changes in the modulation of the expression of the isoforms of different kilodaltons of WT1 in the adaptation process of MCF-7 cells cultured in estrogen depletion, mainly involving the presence of 36-38 kDa WT1 isoform and the loss of the expression of 52-54 Kda WT1 isoform. This behavior is related to the tumor status of the analyzed cell lines, where it is observed that only the positive ER cell lines express the 52-54 kDa WT1 isoform and this is absent in the negative ER cell lines. There are currently few studies where the 36-38 kDa WT1 isoform is analyzed, so it is important to carry out a characterization of the WT1 isoforms in samples of breast cancer patients with various tumor marker status. Finally, we conclude that it is evident that the change of WT1 isoforms during depletion of estrogens in the MCF-7 cells plays a role in adapting to hormonal non-dependence.

## Acknowledgments

This work was supported by the Immunology and Virology Department from Faculty of Biological Sciences, Autonomous University of Nuevo Leon (UANL) and Support Program for Scientific and Technological Research PAICYT-UANL (SA602-18).

## References

1. Ambu R, Vinci L, Gerosa C, Fanni D, Obinu E, Faa A, et al. WT1 expression in the human fetus during development. Eur J Histochem. 2015;59(2):2499.

2. Tatsumi N, Hojo N, Sakamoto H, Inaba R, Moriguchi N, Matsuno K, et al. Identification of a Novel C-Terminal Truncated WT1 Isoform with Antagonistic Effects against Major WT1 Isoforms. PLoS One. 2015;10(6):e0130578.

3. Mayo MW, Wang CY, Drouin SS, Madrid LV, Marshall AF, Reed JC, et al. WT1 modulates apoptosis by transcriptionally upregulating the bcl-2 proto-oncogene. EMBO J. 1999;18(14):3990–4003.

4. Han Y, San-Marina S, Liu J, Minden MD. Transcriptional activation of c-myc proto-oncogene by WT1 protein. Oncogene. 2004;23(41):6933–41.

5. Simpson LA, Burwell EA, Thompson KA, Shahnaz S, Chen AR, Loeb DM. The antiapoptotic gene A1/BFL1 is a WT1 target gene that mediates granulocytic differentiation and resistance to chemotherapy. Blood. 2006;107(12):4695–702.

6. McCarty G, Awad O, Loeb DM. WT1 protein directly regulates expression of vascular endothelial growth factor and is a mediator of tumor response to hypoxia. J Biol Chem. 2011;286(51):43634–43.

7. Bansal H, Seifert T, Bachier C, Rao M, Tomlinson G, Iyer SP, et al. The transcription factor Wilms tumor 1 confers resistance in myeloid leukemia cells against the proapoptotic therapeutic agent TRAIL (tumor necrosis factor alpha-related apoptosis-inducing ligand) by regulating the antiapoptotic protein Bcl-xL. J Biol Chem. 2012;287(39):32875–80.

8. Xu C, Wu C, Xia Y, Zhong Z, Liu X, Xu J, et al. WT1 promotes cell proliferation in non-small cell lung cancer cell lines through up-regulating cyclin D1 and p-pRb in vitro and in vivo. PLoS One. 2013;8(8):e68837.

9. Moffett P, Bruening W, Nakagama H, Bardeesy N, Housman D, Housman DE, et al. Antagonism of WT1 activity by protein self-association. Proc Natl Acad Sci U S A. 1995;92(24):11105–9.

10. Reddy JC, Morris JC, Wang J, English MA, Haber DA, Shi Y, et al. WT1-mediated transcriptional activation is inhibited by dominant negative mutant proteins. J Biol Chem. 1995;270(18):10878–84.

11. Kennedy D, Ramsdale T, Mattick J, Little M. An RNA recognition motif in Wilms’ tumour protein (WT1) revealed by structural modelling. Nat Genet. 1996;12(3):329–31.

12. Rauscher FJ, Morris JF, Tournay OE, Cook DM, Curran T. Binding of the Wilms’ tumor locus zinc finger protein to the EGR-1 consensus sequence. Science. 1990;250(4985):1259–62.

13. Bardeesy N, Pelletier J. Overlapping RNA and DNA binding domains of the wt1 tumor suppressor gene product. Nucleic Acids Res. 1998;26(7):1784–92.

14. Nakagama H, Heinrich G, Pelletier J, Housman DE. Sequence and structural requirements for high-affinity DNA binding by the WT1 gene product. Mol Cell Biol. 1995;15(3):1489–98.

15. Dechsukhum C, Ware JL, Ferreira-Gonzalez A, Wilkinson DS, Garrett CT. Detection of a novel truncated WT1 transcript in human neoplasia. Mol Diagn. 2000;5(2):117–28.

16. Tuna M, Itamochi H. Insulin-like growth factor I regulates the expression of isoforms of Wilms’ tumor 1 gene in breast cancer. Tumori. 2013;99(6):715–22.

17. Haber DA, Sohn RL, Buckler AJ, Pelletier J, Call KM, Housman DE. Alternative splicing and genomic structure of the Wilms tumor gene WT1. Proc Natl Acad Sci U S A. 1991;88(21):9618–22.

18. Bruening W, Pelletier J. A non-AUG translational initiation event generates novel WT1 isoforms. J Biol Chem. 1996;271(15):8646–54.

19. Scharnhorst V, Dekker P, van der Eb AJ, Jochemsen AG. Internal translation initiation generates novel WT1 protein isoforms with distinct biological properties. J Biol Chem. 1999;274(33):23456–62.

20. Dallosso AR, Hancock AL, Brown KW, Williams AC, Jackson S, Malik K. Genomic imprinting at the WT1 gene involves a novel coding transcript (AWT1) that shows deregulation in Wilms’ tumours. Hum Mol Genet. 2004;13(4):405–15.

21. Hossain A, Nixon M, Kuo MT, Saunders GF. N-terminally truncated WT1 protein with oncogenic properties overexpressed in leukemia. J Biol Chem. 2006;281(38):28122–30.

22. Oji Y, Miyoshi S, Maeda H, Hayashi S, Tamaki H, Nakatsuka S, et al. Overexpression of the Wilms’ tumor gene WT1 in de novo lung cancers. Int J Cancer. 2002;100(3):297–303.

23. Oji Y, Yamamoto H, Nomura M, Nakano Y, Ikeba A, Nakatsuka S, et al. Overexpression of the Wilms’ tumor gene WT1 in colorectal adenocarcinoma. Cancer Sci. 2003;94(8):712–7.

24. Oji Y, Nakamori S, Fujikawa M, Nakatsuka S, Yokota A, Tatsumi N, et al. Overexpression of the Wilms’ tumor gene WT1 in pancreatic ductal adenocarcinoma. Cancer Sci. 2004;95(7):583–7.

25. Loeb DM, Evron E, Patel CB, Sharma PM, Niranjan B, Buluwela L, et al. Wilms’ tumor suppressor gene (WT1) is expressed in primary breast tumors despite tumor-specific promoter methylation. Cancer Res. 2001;61(3):921–5.

26. Oji Y, Ogawa H, Tamaki H, Oka Y, Tsuboi A, Kim EH, et al. Expression of the Wilms’ tumor gene WT1 in solid tumors and its involvement in tumor cell growth. Jpn J Cancer Res. 1999;90(2):194–204.

27. Yamauchi T, Negoro E, Lee S, Takai M, Matsuda Y, Takagi K, et al. Detectable Wilms’ tumor-1 transcription at treatment completion is associated with poor prognosis of acute myeloid leukemia: a single institution’s experience. Anticancer Res. 2013;33(8):3335–40.

28. Drakos E, Rassidakis GZ, Tsioli P, Lai R, Jones D, Medeiros LJ. Differential expression of WT1 gene product in non-Hodgkin lymphomas. Appl Immunohistochem Mol Morphol. 2005;13(2):132–7.

29. Silberstein GB, Van Horn K, Strickland P, Roberts CT, Jr., Daniel CW. Altered expression of the WT1 wilms tumor suppressor gene in human breast cancer. Proc Natl Acad Sci U S A. 1997;94(15):8132–7.

30. Miyoshi Y, Ando A, Egawa C, Taguchi T, Tamaki Y, Tamaki H, et al. High expression of Wilms’ tumor suppressor gene predicts poor prognosis in breast cancer patients. Clin Cancer Res. 2002;8(5):1167–71.

31. Oji Y, Miyoshi Y, Kiyotoh E, Koga S, Nakano Y, Ando A, et al. Absence of mutations in the Wilms’ tumor gene WT1 in primary breast cancer. Jpn J Clin Oncol. 2004;34(2):74–7.

32. Zapata-Benavides P, Tuna M, Lopez-Berestein G, Tari AM. Downregulation of Wilms’ tumor 1 protein inhibits breast cancer proliferation. Biochem Biophys Res Commun. 2002;295(4):784–90.

33. Tuna M, Chavez-Reyes A, Tari AM. HER2/neu increases the expression of Wilms’ Tumor 1 (WT1) protein to stimulate S-phase proliferation and inhibit apoptosis in breast cancer cells. Oncogene. 2005;24(9):1648–52.

34. Livak KJ, Schmittgen TD. Analysis of relative gene expression data using real-time quantitative PCR and the 2(-Delta Delta C(T)) Method. Methods. 2001;25(4):402–8.

35. Ghoncheh M, Pournamdar Z, Salehiniya H. Incidence and Mortality and Epidemiology of Breast Cancer in the World. Asian Pac J Cancer Prev. 2016;17(S3):43–6.

36. Dai X, Li T, Bai Z, Yang Y, Liu X, Zhan J, et al. Breast cancer intrinsic subtype classification, clinical use and future trends. Am J Cancer Res. 2015;5(10):2929–43.

37. Russnes HG, Lingjærde OC, Børresen-Dale AL, Caldas C. Breast Cancer Molecular Stratification: From Intrinsic Subtypes to Integrative Clusters. Am J Pathol. 2017;187(10):2152–62.

38. Wang L, Wang ZY. The Wilms’ tumor suppressor WT1 induces estrogen-independent growth and anti-estrogen insensitivity in ER-positive breast cancer MCF7 cells. Oncol Rep. 2010;23(4):1109–17.

39. Hutcheson IR, Knowlden JM, Madden TA, Barrow D, Gee JM, Wakeling AE, et al. Oestrogen receptor-mediated modulation of the EGFR/MAPK pathway in tamoxifen-resistant MCF-7 cells. Breast Cancer Res Treat. 2003;81(1):81–93.

40. Yang L, Han Y, Suarez Saiz F, Minden MD. A tumor suppressor and oncogene: the WT1 story. Leukemia. 2007;21(5):868–76.

41. Oji Y, Inohara H, Nakazawa M, Nakano Y, Akahani S, Nakatsuka S, et al. Overexpression of the Wilms’ tumor gene WT1 in head and neck squamous cell carcinoma. Cancer Sci. 2003;94(6):523–9.

42. Ueda T, Oji Y, Naka N, Nakano Y, Takahashi E, Koga S, et al. Overexpression of the Wilms’ tumor gene WT1 in human bone and soft-tissue sarcomas. Cancer Sci. 2003;94(3):271–6.

43. Burwell EA, McCarty GP, Simpson LA, Thompson KA, Loeb DM. Isoforms of Wilms’ tumor suppressor gene (WT1) have distinct effects on mammary epithelial cells. Oncogene. 2007;26(23):3423–30.

44. Siehl JM, Reinwald M, Heufelder K, Menssen HD, Keilholz U, Thiel E. Expression of Wilms’ tumor gene 1 at different stages of acute myeloid leukemia and analysis of its major splice variants. Ann Hematol. 2004;83(12):745–50.

45. Gillmore R, Xue SA, Holler A, Kaeda J, Hadjiminas D, Healy V, et al. Detection of Wilms’ tumor antigen--specific CTL in tumor-draining lymph nodes of patients with early breast cancer. Clin Cancer Res. 2006;12(1):34–42.

46. Ito K, Oji Y, Tatsumi N, Shimizu S, Kanai Y, Nakazawa T, et al. Antiapoptotic function of 17AA(+)WT1 (Wilms’ tumor gene) isoforms on the intrinsic apoptosis pathway. Oncogene. 2006;25(30):4217–29.

47. Tatsumi N, Oji Y, Tsuji N, Tsuda A, Higashio M, Aoyagi S, et al. Wilms’ tumor gene WT1-shRNA as a potent apoptosis-inducing agent for solid tumors. Int J Oncol. 2008;32(3):701–11.

48. Graidist P, Nawakhanitworakul R, Saekoo J, Dechsukhum C, Fujise K. Anti-apoptotic function of T-KTS+, T-KTS-, WT1+/+ and WT1+/− isoforms in breast Cancer. 2010;4(5):711.

49. Jomgeow T, Oji Y, Tsuji N, Ikeda Y, Ito K, Tsuda A, et al. Wilms’ tumor gene WT1 17AA(−)/KTS(−) isoform induces morphological changes and promotes cell migration and invasion in vitro. Cancer Sci. 2006;97(4):259–70.

50. Inoue K, Tamaki H, Ogawa H, Oka Y, Soma T, Tatekawa T, et al. Wilms’ tumor gene (WT1) competes with differentiation-inducing signal in hematopoietic progenitor cells. Blood. 1998;91(8):2969–76.

51. Tsuboi A, Oka Y, Ogawa H, Elisseeva OA, Tamaki H, Oji Y, et al. Constitutive expression of the Wilms’ tumor gene WT1 inhibits the differentiation of myeloid progenitor cells but promotes their proliferation in response to granulocyte-colony stimulating factor (G-CSF). Leuk Res. 1999;23(5):499–505.

52. Cheema SK, Mishra SK, Rangnekar VM, Tari AM, Kumar R, Lopez-Berestein G. Par-4 transcriptionally regulates Bcl-2 through a WT1-binding site on the bcl-2 promoter. J Biol Chem. 2003;278(22):19995–20005.

53. Maheswaran S, Park S, Bernard A, Morris JF, Rauscher FJ, 3rd, Hill DE, et al. Physical and functional interaction between WT1 and p53 proteins. Proc Natl Acad Sci U S A. 1993;90(11):5100–4.

54. Richard DJ, Schumacher V, Royer-Pokora B, Roberts SG. Par4 is a coactivator for a splice isoform-specific transcriptional activation domain in WT1. Genes Dev. 2001;15(3):328–39.

